# Power, false discovery rate and Winner’s Curse in eQTL studies

**DOI:** 10.1101/209171

**Authors:** Qin Qin Huang, Scott C. Ritchie, Marta Brozynska, Michael Inouye

## Abstract

Investigation of the genetic architecture of gene expression traits has aided interpretation of disease and trait-associated genetic variants, however key aspects of expression quantitative trait (eQTL) study design and analysis remain understudied. We used extensive, empirically-driven simulations to explore eQTL study design and the performance of various analysis strategies. Across multiple testing correction methods, false discoveries of genes with eQTLs (eGenes) were substantially inflated when false discovery rate (FDR) control was applied to all tests, and only appropriately controlled using hierarchical procedures. All multiple testing correction procedures had low power and inflated FDR for eGenes whose causal SNPs had small allele frequencies using small sample sizes (e.g. frequency <10% in 100 samples), indicating that even moderately low frequency eQTL SNPs (eSNPs) in these studies are enriched for false discoveries. In scenarios with ≥80% power, the top eSNP was the true simulated eSNP 90% of the time, but substantially less frequently for very common eSNPs (minor allele frequencies >25%). Overestimation of eQTL effect sizes, so-called “Winner’s Curse”, was common in low and moderate power settings. To address this, we developed a bootstrap method (BootstrapQTL) which led to more accurate effect size estimation. These insights provide a foundation for future eQTL studies, especially those with sampling constraints and subtly different conditions.

## Introduction

Genome-wide association studies (GWAS) have identified thousands of genetic variants associated with complex phenotypes^1^ and the vast majority of genome-wide significant SNPs are located in non-coding region^2^, making interpretation challenging. Integration of gene expression and genetic variation is a ubiquitous approach for uncovering genetic regulatory effects and their ramifications for pathways relevant to human diseases and traits^3,4,5,6^, and indeed trait-associated SNPs have been found to be enriched for expression quantitative trait loci (eQTL) effects^7^.

Yet, while eQTL analysis has become a focus of functional genomics, the lack of a strong evidence base for eQTL study design leaves fundamental questions unanswered. In particular, while more and more eQTLs reach statistical significance, the true proportion of false discoveries and the accuracy of their effect size estimates have not yet been well characterised. A seminal early study compared multiple testing correction methods for detecting eQTLs (including Bonferroni correction, false discovery rate control and permutation) using HapMap data, however estimates of false discovery rate (FDR) and sensitivity are not possible without knowledge of all true eQTLs in the data^8^. Previous eQTL simulations are typically part of new methodologies, yet these simulations have been limited in their reflection of real data. Genotype data have typically been simulated with a narrow minor allele frequency (MAF) range assuming Hardy-Weinberg equilibrium (e.g. MAF 30% in Ref ^9^, 5% and 20% in Ref ^10^, 40% in Ref ^11^), thus they have not captured realistic patterns of genetic variation, especially linkage disequilibrium (LD) complexity. Furthermore, MAFs at 1% or greater are typically utilized for eQTL analysis (**Table S1**). Others have simulated only a fixed sample size^11,12,13^. Typically, eQTL studies have sample sizes of 50 to 1,000, with the accessibility of the tissue, cell-type or condition a major determining factor (**Table S1**). A recent *trans*-eQTL study performed in whole blood had a size of 5,257 samples^6^ and a study combined data for 2,116 whole blood samples to identify context-specific eQTLs^14^. Perhaps the exemplar multiple human tissue resource, the Genotype-Tissue Expression (GTEx) project^15^, comprises 44 tissues with a sample size range of 70–361 in its V6p data release^16^.

While studies have generally converged on linear regression or linear mixed models for eQTL detection, the multiple testing correction approach is still a source of substantial variability among studies. Various approaches are available for minimizing type I errors. Often criticised as too conservative, particularly with complex LD patterns, the Bonferroni correction aims to control the familywise error rate (the probability of making any type I error) by setting the significance level at α/N, where α is the desired significance level (0.05 conventionally) and N is the number of tests. FDR-controlling procedures, which aim to control the expected proportion of false discoveries among all rejected null hypotheses, are generally considered to provide a better balance between false positives and false negatives. Benjamini and Hochberg (BH) proposed a procedure^17^ assuming each statistical test is independent, which is not the case due to LD. Benjamini and Yekutieli (BY) modified the FDR procedure to one which, while more conservative, accommodates correlation structure between statistical tests^18^. The q-value FDR-controlling approach from Storey and Tibshirani (ST) estimates the proportion of hypotheses that are truly null (π_0_), while the BH procedure assumes π_0_ = 1 which makes ST less conservative than the BH procedure^19^.

Other approaches have been proposed to deal with multiple testing specifically for eQTL studies. Locus-restricted permutation testing is widely used to obtain empirical null distributions. To achieve this, sample labels are randomly shuffled while keeping genotype data constant, with association tests performed at each permutation step. For each gene, the best SNP association at each permutation is kept to generate an empirical null distribution of minimum p-values, from which permutation test p-values are calculated for each *cis*-SNP. Thousands of permutations are required to achieve accurate results, thus there is a high computational cost. Approximations have been investigated for calculating permutation p-values, such as those in FastQTL^20^ and MVN^21^. For example, FastQTL provides an option to approximate the tail of the empirical null distributions of p-values using a beta distribution thereby reducing the number of permutations required^20^. In addition to permutation tests, eigenMT proposed by Davis *et al.*^22^ adjusts p-values in shorter time. The number of independent tests (typically SNPs) for each gene is estimated by eigenMT using a genotype correlation matrix, then a Bonferroni procedure is applied^22^. Both FastQTL and eigenMT account for LD structure among local variants. Recently, hierarchical procedures, such as TreeQTL^23^, have been proposed, which first control for multiple testing of variants at each gene, before controlling for multiple testing across all genes. Taken together, with many correction methods available, it is not clear which method is optimal for eQTL mapping nor what their respective performances are for genetic variants with difference characteristics (allele frequency, effect size, etc).

Effect size estimation for eQTLs represents a more complex and less explored problem, yet its importance is increasing as comparison of eQTLs across tissues, experimental conditions, and meta-analyses become more common. Furthermore, prediction of tissue-specific gene expression from genotypes, for example using the tool PrediXcan^24^, is critically dependent on effect size estimation, particularly *cis*-eQTL effect sizes obtained from analyses of GTEx and other studies. Conversely, a method which predicts genotypes at eQTL SNPs (eSNPs) based on measured gene expression levels has also been proposed^25^.

A well-recognised phenomenon in GWAS is “Winner’s Curse”^26,27^, an ascertainment bias where the true genetic effect is smaller than its estimate within the discovery cohort, a problem which is accentuated when power is low. Methods have been proposed to correct this upward bias, such as those based on likelihood^28^ and resampling^29^, however none have been tailored to eQTLs. Winner’s Curse has also been reported in eQTL studies^30^ but its presence and methods for its correction have not been systematically evaluated.

Here we used extensive simulations of realistic LD patterns of human genetic variation and matched gene expression to investigate how various scenarios, including different sample sizes, allele frequencies and genetic effect sizes, influence statistical power and FDR (Fig. 1). In each scenario, we randomly selected SNPs as true causal *cis*-eQTLs, each associated with expression levels of a target gene. We performed eQTL mapping and evaluated a variety of multiple testing correction methods, used both individually and hierarchically, under each scenario. We next investigated the accuracy of genetic effect size estimation across scenarios, the effect of the Winner’s Curse, and how bias was affected by study power. Finally, we evaluated the accuracy of a variety of eQTL effect size estimation procedures.

**Figure 1.**
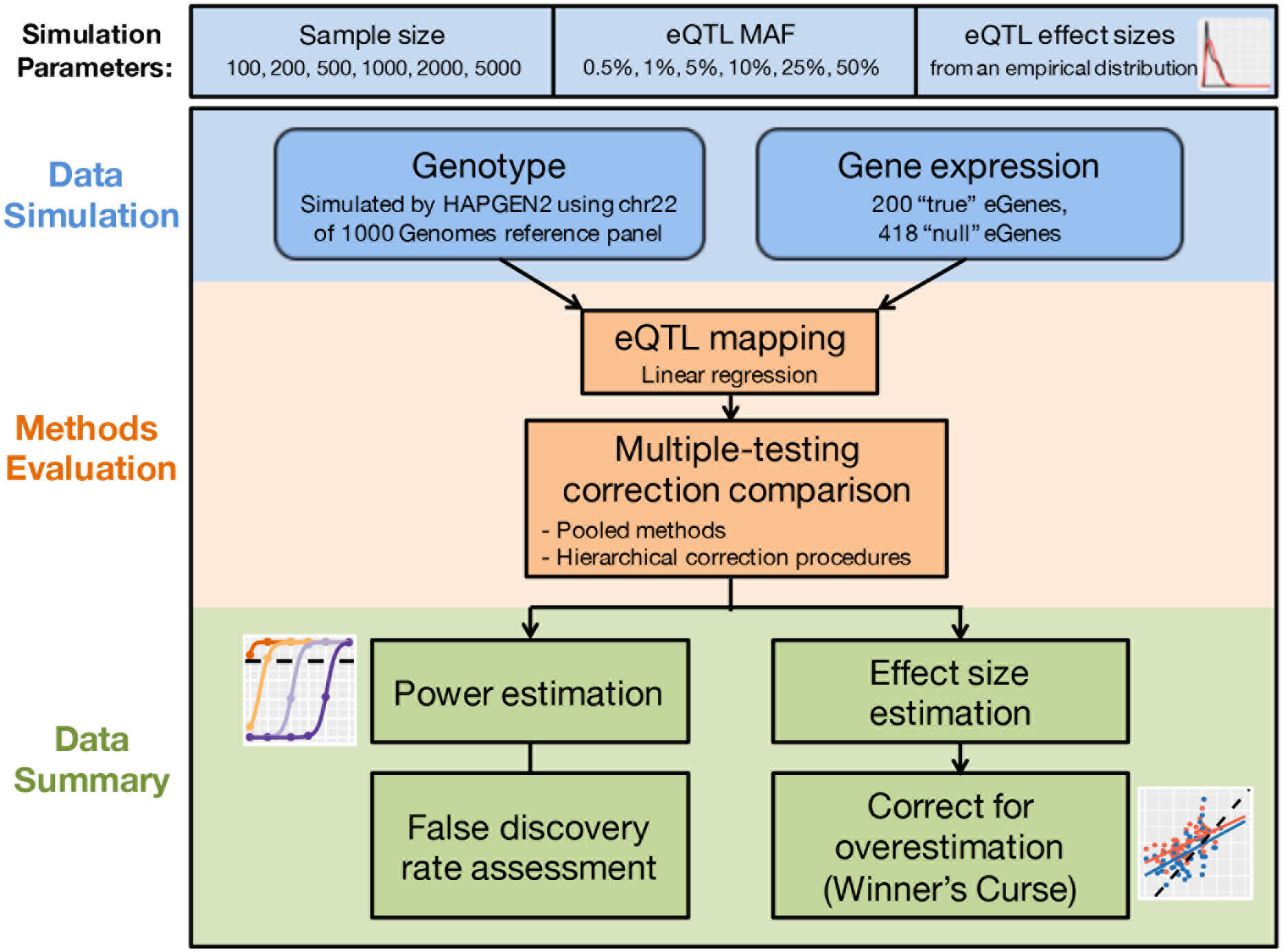
Flowchart our eQTL simulation study.

## Results

### Simulation of *cis*-eQTL data

To assess the power, FDR, and effect size estimation of eQTL studies based on different parameters, we simulated 36 scenarios with combinations of six sample sizes (N=100, 200, 500, 1000, 2000, and 5000) and six true minor allele frequencies of eSNPs (MAF=0.5%, 1%, 5%, 10%, 25%, and 50%). Realistic LD patterns were simulated using HAPGEN2^31^ with chromosome 22 of the 1000 Genomes Project phase 3 data^32^ as reference. In each scenario, 618 gene expression traits were simulated, among which 200 were under genetic regulation (true eGenes). Each true eGene was simulated to be regulated by one *cis*-eQTL with a genetic effect size randomly drawn from an empirical distribution based on eQTL analysis of a real dataset^33,34^.

For each gene, all SNPs located within 1Mb of the transcription start site (TSS) were tested for association using linear regression models through Matrix eQTL^35^. We mapped *cis-* eQTLs for the 36 scenarios separately and evaluated different multiple testing correction methods. **Figure 1** illustrates the workflow of our eQTL simulations and methods evaluation. We used Bonferroni, FDR-controlling procedures, permutation approaches, and eigenMT to correct for multiple testing. The Bonferroni and FDR procedures were applied alone to all hypotheses (pooled method) and were also used in combination via a hierarchical correction procedure (**Methods**). We repeated the simulation for each scenario 100 times and calculated the sensitivity and FDR of each multiple testing correction method based on all simulations.

### Power and false discovery rate between scenarios and multiple testing correction procedures

We first assessed the variability in sensitivity and FDR for the various multiple testing correction methods for eGene detection across simulation scenarios. A significant eGene was considered a true positive if: (1) it was among the 200 true eGenes simulated, and (2) the simulated causal eSNP for that eGene was among the significant eSNPs, or a significant eSNP was in high LD with the causal eSNP (r^2^ ≥0.8). For each multiple testing correction method, sensitivity, or true positive rate (TPR), was calculated as the proportion of simulated true eGenes correctly identified as true positives. Conversely, the FDR was calculated as the proportion of false positives in significant eGenes identified across all 100 simulations. We did not calculate FDR for a scenario if no eGenes were significant in all simulations.

We evaluated multiple testing correction methods in two ways; first, applied across all SNP–gene hypothesis tests (hereby “pooled methods”), and second, in combinations in a hierarchical approach in which SNP–gene hypothesis tests were partitioned into groups by the gene being tested (hereby “hierarchical correction procedures”) ^36^. In the case of hierarchical correction procedures, the multiple hypothesis tests of eGenes were controlled (Step 2, global correction) based on the multiple testing adjusted statistics (Step 1, local correction) of each gene’s best association, then SNPs significantly associated with the significant eGenes were identified based on the locally corrected p-value corresponding to the threshold of 0.05 after global correction (Step 3, **Methods**).

FDR-controlling procedures applied to all hypotheses (pooled FDR methods) failed to control the FDR of eGenes in nearly all scenarios, and FDR increased with statistical power (**Fig. S1**). We applied three FDR-controlling procedures to all hypotheses: the Storey-Tibshirani (ST)^19^, Benjamini-Hochberg (BH)^17^, and Benjamini-Yekutieli (BY)^18^ procedures. The ST and BH procedures failed to control FDR at the desired level of 0.05 in majority of the scenarios, and FDR increased with sample size, reaching more than 0.6 under scenarios with sample sizes of 2,000 or 5,000 and true eSNP MAFs ≥25% (**Fig. S1A**). The BY procedure was the most conservative method among pooled FDR procedures but still had inflated FDR under scenarios with large sample sizes (≥1,000) and true eSNP MAFs ≥25%. As expected, a pooled Bonferroni correction had very low FDR values in most scenarios, with the lowest sensitivity across MAFs and sample sizes (**Fig. S1**). However, even pooled Bonferroni correction failed to control FDR of rare variant eQTLs (MAF ≤1%) in scenarios with <1,000 samples. Overall, we observed inflated rates of false positive eGenes for all pooled FDR methods.

In contrast to pooled methods, we observed better calibrated FDR for hierarchical multiple testing correction procedures, except in scenarios with low statistical power (**Fig. 2A**, **Fig. S2**). We compared ST, BH, BY, Bonferroni, eigenMT, and three permutation approaches (discussed in a later paragraph) for adjusting the *cis*-SNP p-values for each simulated gene (local correction), combined with a comparison of the ST, BH, BY, and Bonferroni correction for adjusting the subsequent minimum adjusted p-value across all genes (global correction).

**Figure 2.**
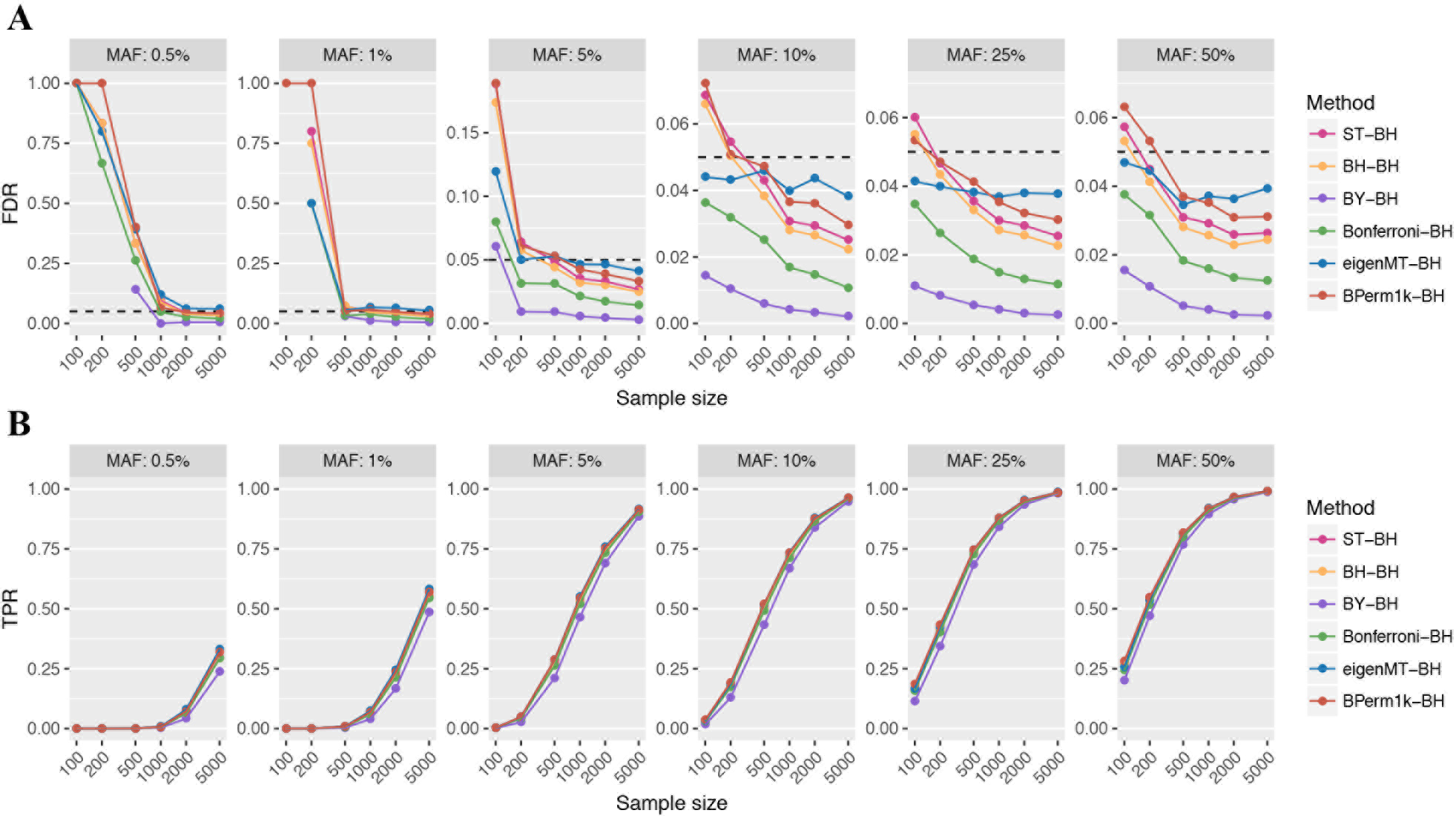
False discovery rate (FDR) and sensitivity of selected hierarchical multiple testing correction methods. Comparison of the FDR (**A**) and sensitivity (**B**) of six methods (different colours) for controlling multiple testing of SNPs at each gene (local correction), with Benjamini-Hochberg (BH) used to control for multiple testing across all genes (global correction). The six methods compared were Storey-Tibshirani (ST), Benjamini-Hochberg (BH), Benjamini-Yekutieli (BY), Bonferroni correction, eigenMT, and permutation tests based on beta approximation (BPerm1k). Comparison of all combinations of multiple testing correction methods for hierarchical correction are shown in **Fig. S2** and **Fig. S3**. Application of BH in the global correction step had the best sensitivity for all methods used in the local correction step of any hierarchical correction procedures. Each dot represents one scenario and plots show different minor allele frequencies (MAFs) of the simulated causal eSNPs. The dashed horizontal line indicates the desired FDR level of 5%. Scenarios where no significant eGenes were identified are not shown in panel A.

We observed lower sensitivity as well as lower FDR than ST and BH when applying BY and Bonferroni to correct across genes, regardless of which multiple testing correction method was used for local correction (**Fig. S2**, **Fig. S3**). ST and BH global correction had identical performance, except when permutation tests were used as local correction method, where ST had higher FDR than BH and often had FDR slightly higher than 5% (**Fig. S2**, **Fig. S3**). We therefore subsequently focused on the BH procedure to control for multiple testing across genes in hierarchical correction procedures.

We compared three different permutation approaches to correct for multiple testing at each gene: (1) using exact permutation test p-values from 1,000 permutations (Perm1k-BH), (2) using p-values obtained from beta distribution approximation of each null distribution’s tail after 1,000 permutations (BPerm1k-BH), and (3) using beta approximation under an adaptive scheme where a minimum of 100 and a maximum of 10,000 permutations were performed for each gene based on the significance level of this gene (APerm10k-BH). Due to the prohibitive computational time required to run Perm1k and APerm10k, we ran 10 simulations rather than 100 to compare the three permutation approaches. Perm1k-BH had lower sensitivity than the other two permutation approaches in scenarios with low detection power and it also had a higher FDR (**Fig. S4**). BPerm1k and APerm10k had similar performance, indicating 1000 permutations were sufficient to obtain an accurate approximation of the p-value null distribution tail. We therefore used BPerm1k-BH as a representative of permutation approaches to compare with other multiple testing correction methods.

Amongst the hierarchical correction methods with BH as global correction, BY adjustment of multiple SNPs (BY-BH) had the most conservative FDR among all methods, more so than Bonferroni-BH due to BY’s heavier correction for the lowest p-values; however, this came at the expense of lower sensitivity (**Fig. 2**). Besides BY-BH, other methods did not show a notable difference in sensitivity. Perhaps surprisingly, Bonferroni-BH maintained a comparable sensitivity to other methods while having an FDR well below 0.05. In terms of calibration, eigenMT-BH had an FDR closest to 0.05 and was relatively stable with respect to sample size, whereas other methods showed an inverse relationship between FDR and sample size. In the Discussion, we explore the trade-offs of FDR calibration versus minimization for a given power. Below, we utilise the eigenMT-BH procedure to illustrate the ramification of our findings for eQTL study design, while also noting that design differences between Bonferroni-BH and eigenMT-BH would be minor.

Across all effect sizes and using the eigenMT-BH procedure (**Fig. 2**), it was apparent that (i) eSNPs with ≤0.5% MAF and ≤1% MAF that were detected with <1000 and <500 samples, respectively, were likely to be false discoveries, (ii) for studies with 100 samples, a MAF threshold of 10% is necessary to control FDR at ≤5% irrespective of hierarchical multiple testing procedure. In varying the eSNP effect size (0.25, 0.5, 1.0, or 1.5 s.d. gene expression per allele), we found that sample sizes up to 200 (quite common in the eQTL literature) only reached 80% power for eQTLs of ≥5% MAF and effect size 1.5 s.d. per allele or for eQTLs of MAF 50% and effect size of approximately ≥0.6 s.d. per allele (**Fig. 3**). The maximum sample size of 5,000 in our simulations still did not reach 80% power to detect eQTLs with effect size of 0.25 s.d. per allele and 5% MAF. When sample sizes were >1,000 and MAF >10%, eQTLs with effect size of 0.25 s.d. per allele could be detected at power 80%. Studies of 100 samples were underpowered unless eQTLs were moderately common (at least ~25% MAF) and of large effect size (≥1.0 s.d. per allele).

**Figure 3.**
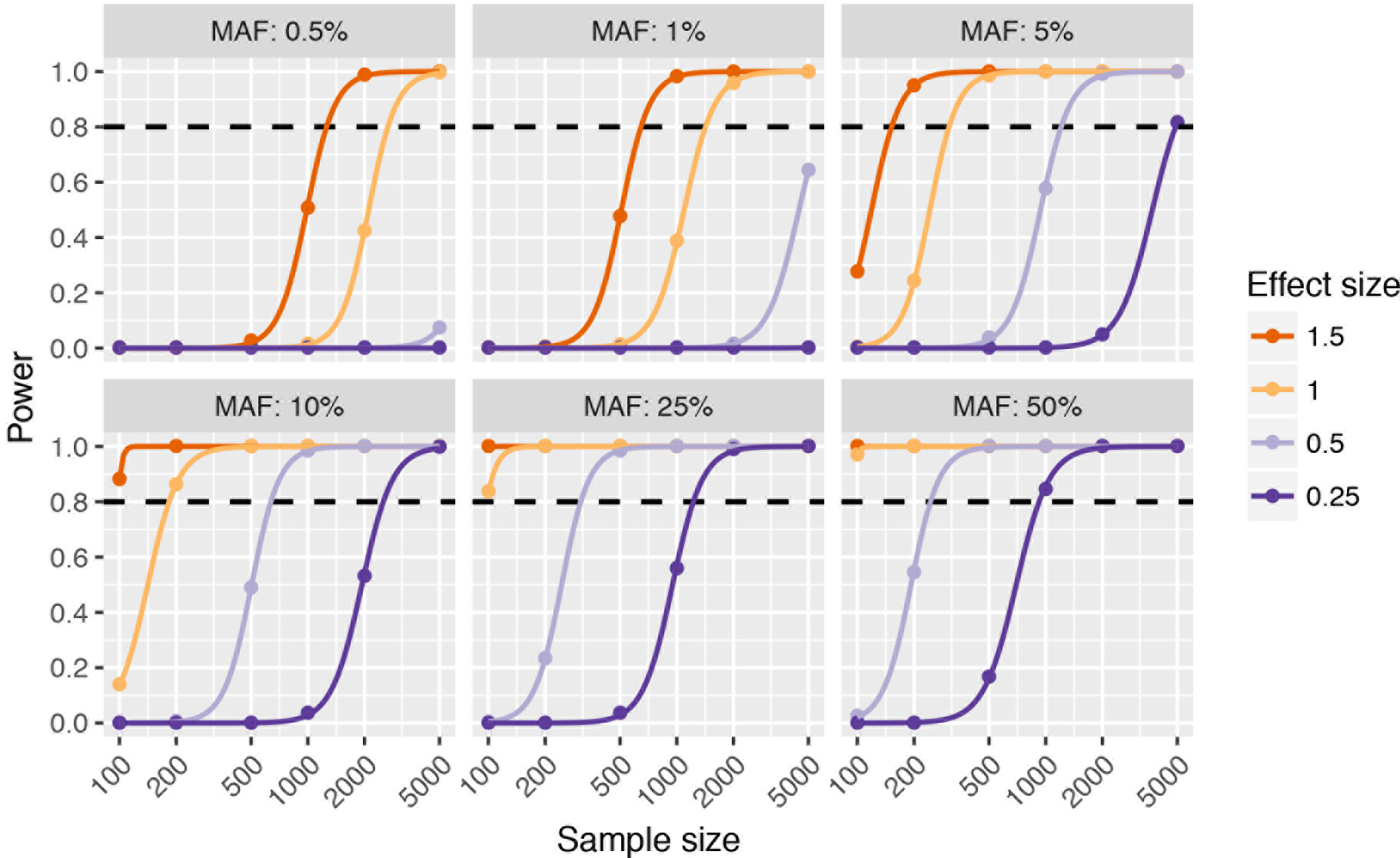
Power and eQTL effect size. A constant genetic effect size (0.25, 0.5, 1.0, or, 1.5 s.d. gene expression per allele) was simulated in each scenario. Plots represents different minor allele frequencies (MAFs) of the simulated true eSNPs. Sample size increases from left to right on x-axes. The estimated statistical power for eGene detection from 100 simulations is shown on y-axes. A hierarchical correction procedure using eigenMT for local correction and BH for global correction (eigenMT-BH) was used to correct for multiple testing. The dashed horizontal line indicates sufficient statistical power (0.8).

### Identification of the simulated causal eSNP

When hierarchical multiple testing correction procedures had calibrated FDR for eGenes, we observed multiple significant eSNPs at each true positive eGene (**Fig. S5**) despite simulating only one causal eSNP for each true eGene, as would be expected given LD. The number of SNPs significantly associated with a true eGene increased with both sample size and true eSNP MAF, with >1,000 significant eSNPs identified per eGene on average in the scenario with the largest sample size (N=5,000), true eSNP MAF (50%), and eQTL effect size (1.5 s.d. per allele) (**Fig. S5**).

Many studies focus on the eSNP with the strongest association (lowest p-value) with each eGene (top eSNP) when performing downstream analyses, such as enrichment analysis or effect size estimation^14,16^. In our simulations, we found that while the power to detect the presence of an eQTL increased with increasing MAF, the probability that the true causal eSNP was the top eSNP declined (**Fig. 4A**). However, holding MAF constant and increasing study power (increasing sample size and effect size) resulted in increasing probability to detect the true causal eSNP (**Fig. 4A**). In scenarios with at least 1% power to detect an eQTL, top eSNPs with MAF 0.5% were nearly always the true causal eSNP. Given the critical role of LD in fine-mapping, we confirmed our observations were due to a positive relationship between an eSNPs, MAF and the amount of local LD (**Fig. 4B**). For top eSNPs that were not true causal eSNPs, 83% were in high LD (r^2^ ≥0.8) with the true causal eSNPs (**Fig. S6**). Overall, for studies with 80% power to detect a given eQTL of MAF ≤25%, the top eSNP was the true causal eSNP 90% of the time.

**Figure 4.**
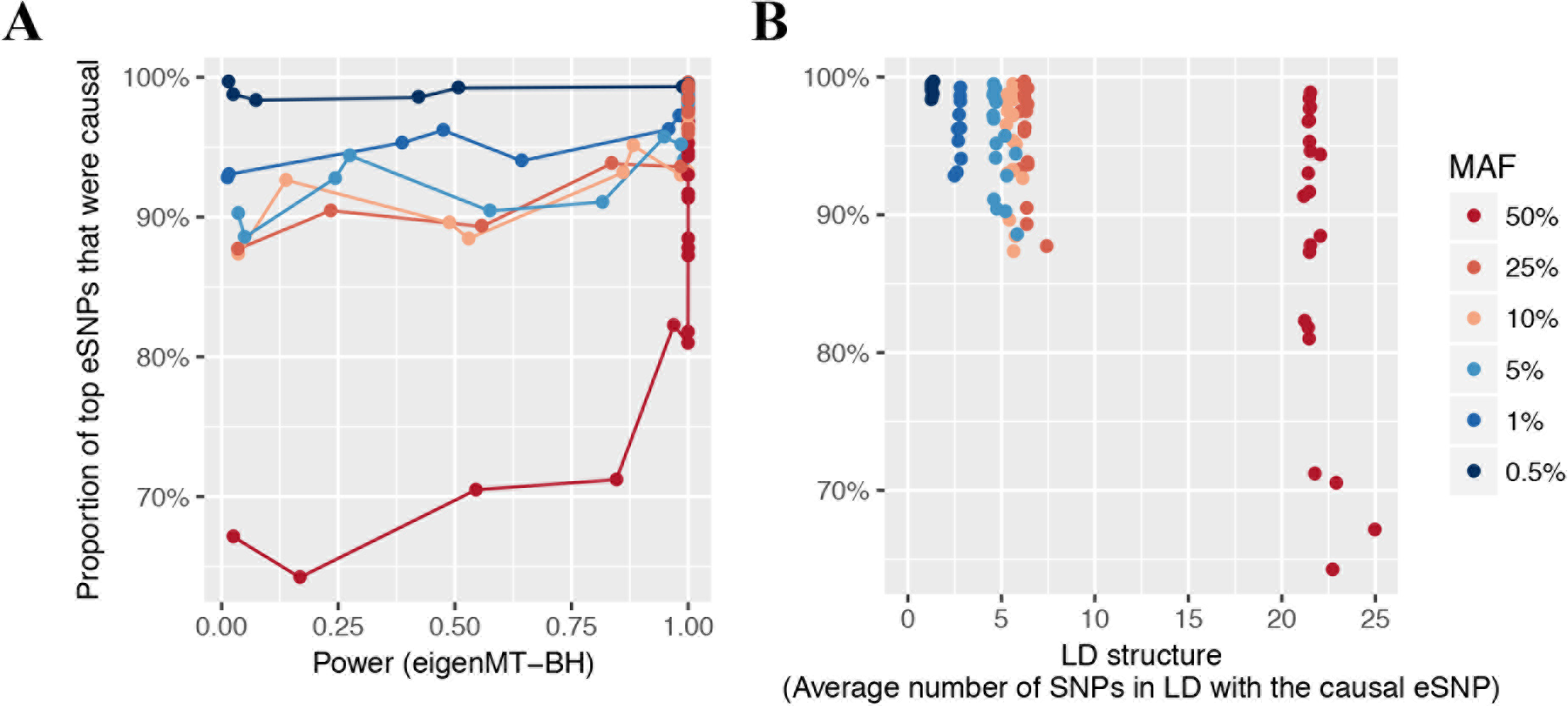
Identification of true causal eSNPs. In each scenario, the 200 causal eSNPs have the same effect size in addition to minor allele frequency (MAF). For significant true eGenes, the proportion of top eSNPs (minimum p-value) that were true causal eSNPs is shown (y-axes) for either (*A*) the power to detect eQTLs of the scenario, or (*B*) the amount of linkage disequilibrium (LD) for true causal eSNPs, i.e. the average number of SNPs within 1Mb and in moderate LD (r^2^ ≥0.5) with the causal eSNP. Scenarios are coloured according to true eSNP minor allele frequency (MAF). Only scenarios with power ≥0.01 are shown. A hierarchical correction procedure using eigenMT for local correction and BH for global correction (eigenMT-BH) was used to identify true positive eGenes.

### Winner’s Curse in eQTL effect size estimation

To systematically evaluate the effect of Winner’s Curse in eQTL studies, we compared beta coefficients obtained from the Matrix eQTL linear regression models for the top eSNP of each true positive eGene (the “naïve estimator”) to their simulated true effect sizes. We observed that median error of the naïve estimator increased as study power decreased, as expected, and also that the naïve estimator consistently overestimated the true effect size with overestimation increasing as power to detect an eQTL decreased (**Fig. 5**, **Fig. S7**).

**Figure 5.**
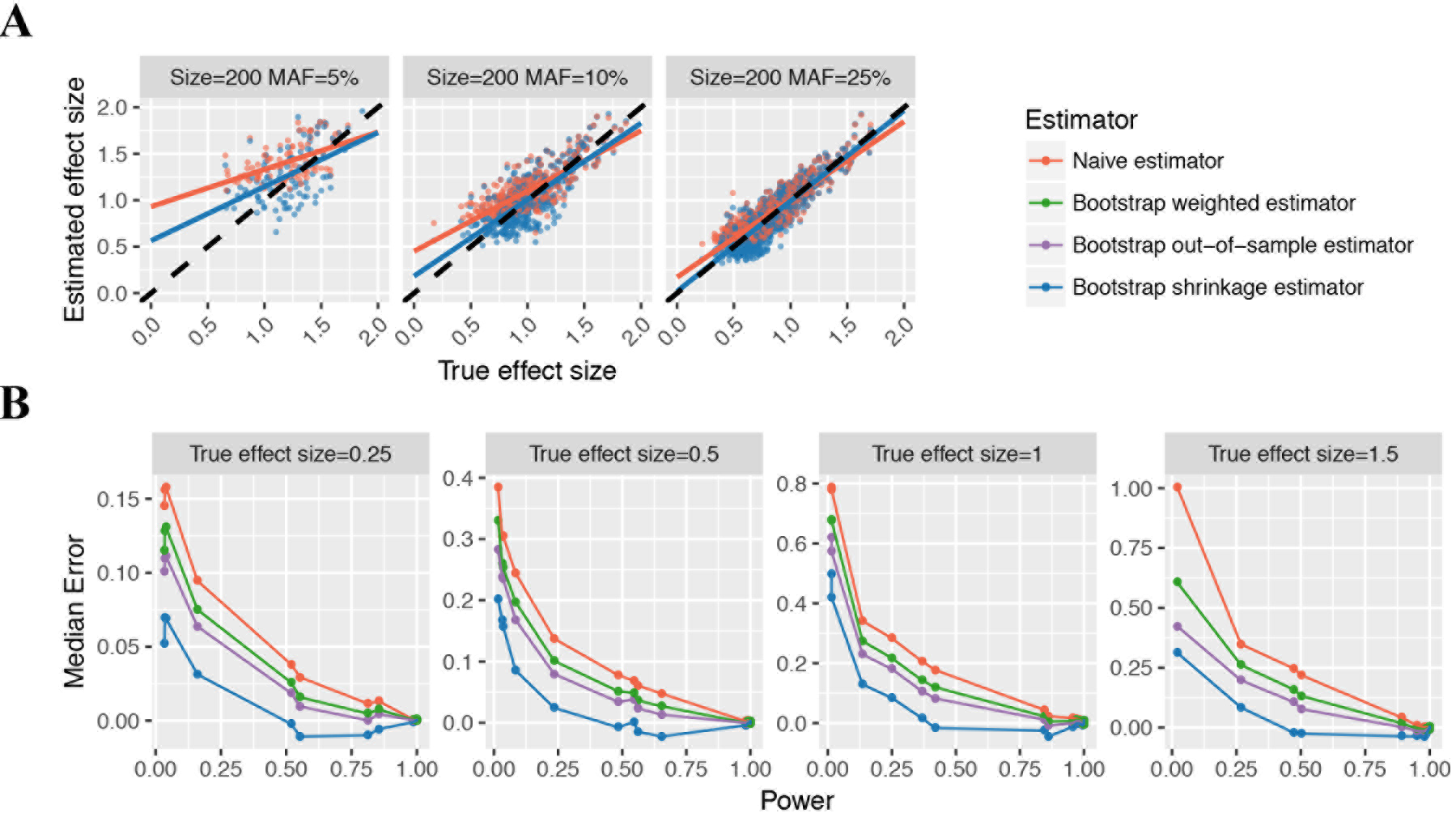
Winner’s Curse in eQTL effect size estimation and correction by bootstrap method. (**A**) shows the phenomenon of Winner’s Curse by three examples: scenarios where the sample size is 200 and the minor allele frequencies (MAFs) of causal eSNPs are 5%, 10%, and 25%. Each dot represents one true positive eGene from ten simulations of the scenario. Plots compare the estimated effect size (y-axes) of the top SNP of each true positive eGene to the true effect size (x-axes) of the simulated causal eSNP. Red points show the naïve estimator (beta coefficient from liner regression) and blue points show the bootstrap shrinkage estimator, which was the best estimator (see panels **B**). Red (or blue) lines are linear regression fit of the naïve estimator (or the bootstrap estimator) on the simulated effect size for the true positive eGenes. Black dashed lines in panel **A** indicate where the estimated effect size equals to the true value. (**B**) shows the median error (difference between estimated and true effect size) for all estimators across 10 simulations of scenarios where a constant true effect size (0.25, 0.5, 1, or, 1.5 s.d. gene expression per allele) was simulated. A hierarchical correction procedure using eigenMT for local correction and BH for global correction (eigenMT-BH) was used to correct for multiple testing.

To address this, we investigated various methods for re-estimating effect sizes. Methods have been proposed to correct for Winner’s Curse in GWAS^28,29^, but to our knowledge, no method has yet been designed for bias correction in eQTL studies. We adapted a bootstrap resampling method^37^ for eQTL studies and compared three bootstrap estimators (a shrinkage estimator, an out-of-sample estimator, and a weighted estimator, see **Methods**) to determine the best approach for adjusting for Winner’s Curse. All three bootstrap estimators had more accurate effect size estimates (smaller mean squared error and median error closer to 0) than the naïve estimator when power of eQTL detection was low to moderate (**Fig. 5B**, **Fig. S8**). Amongst the three bootstrap estimators, the shrinkage estimator was closest to the true effect size overall and across all study powers. In scenarios with high power for eQTL detection, Winner’s Curse was not apparent, and the bootstrap shrinkage estimator and naïve estimator had similar estimates (**Fig. S9**). The bootstrap method for eQTL studies is freely available at https://github.com/InouyeLab/BootstrapQTL.

## Discussion

In this study, we have utilized extensive, realistic simulations of eQTL data to investigate fundamental questions in eQTL study design relating to power, FDR and effect size estimation. The most commonly used MAF cut-offs in recent eQTL studies are 1% or 5% (**Table S1**). For instance, GTEx restricted the association tests to SNPs with minor allele count ≥10 in the tissue analysed, the corresponding MAF being 7% and 1.4%, in the minimum (70) and the maximum (361) sample size, respectively^16^. In our simulations, we found that eQTLs with a small MAF identified in low sample sizes were highly likely to be false positives, regardless of which multiple testing correction strategy was used (**Fig. 2**). Based on above, when 100, 200, and 500 samples are available (typical in eQTL studies), we recommend a MAF cut-off at 10%, 5% and 1%, respectively. Many studies listed in **Table S1** had a lower MAF cut-off than recommended. Detecting rare eQTLs with MAF 0.5% is possible in ≥2000 samples, but even 5,000 samples cannot provide sufficient power unless the eQTL effect size is extremely high: 1.5 s.d. gene expression per allele dosage (**Fig. 2**, **Fig. 3**).

Recent eQTL studies have used pooled FDR methods to correct for multiple testing^38,39,40^. Here, we show that pooled methods are inappropriate for eQTL studies, as they give inflated (sometimes substantially) FDR which worsen as sample size or eSNP MAF increases. This suggests that many eQTLs identified in these studies may be false positives. Hierarchical multiple testing correction procedures had substantially better calibrated FDR. A hierarchical approach of permutation as local correction method followed by ST global adjustment is commonly used in eQTL studies (e.g. by GTEx^16^). When permutation was used as a local correction method, ST often had FDR slightly higher than the desired level in our simulations, while use of BH instead would have better calibrated FDR. Notably, ST and BH adjustment of multiple genes after correction for multiple local SNPs at each gene using other methods except permutation tests had identical results, therefore we recommend using BH to adjust across genes rather than ST.

Most hierarchical procedures had nearly identical sensitivity when BH was used to correct for multiple testing across genes, thus FDR was a differentiating factor. Here, when studies were appropriately powered, eigenMT-BH was the most closely calibrated approach for controlling FDR at 5%, and it had the least variable FDR across different sample sizes. On the other hand, Bonferroni-BH had the smallest FDR with negligibly lower sensitivity. The trade-offs between the use of Bonferroni-BH versus eigenMT-BH are best considered in the context of the specific study. Statistically, calibration is perhaps the deciding factor; if the analysis is intended to guide time-consuming experimental follow-up of specific eQTLs then it may be preferable to minimise FDR for a given detection power.

After eGene detection, identification of the causal eSNP among the significant eSNPs with high LD remains a challenge. Interestingly, we found that the simulated causal eSNP was the most significant eSNP approximately 90% of the time. When the causal variant was not the top variant, ~80% of the time the causal eSNP was in high LD (r^2^ ≥0.8). However, our simulations were simplified to assume that each eGene was regulated by only one casual eSNP, despite observations that gene expression can also be affected by multiple independent genetic variants. 41.2% of protein-coding genes across all 44 tissues showed multiple independent *cis*-eQTLs from the GTEx V6p release^16^. Similarly, a separate study found 26.8% of gene probes had multiple independent eSNPs^41^.

Winner’s Curse in eQTL effect size estimation must be taken into account when comparing effect sizes from different tissue types or conditions, estimating replication sample size, or constructing predictive models. For example, a recent study compared *cis*-eQTL effects between blood samples (N=1,240 samples) and four other tissues (N<85 samples), identifying >2,000 probes with *cis*-eQTL associations that were tissue-dependent, and nearly half were with the same eSNP but with a different effect size^41^. This may be an artefact of Winner’s Curse. To address eQTL effect over-estimation, we have presented a bootstrap method and tool for re-estimation which should enable more accurate eQTL comparisons as well as predictive genetic models for gene expression for less accessible tissues, cell types, conditions or other situations where power is limited.

The investigation of the genetic component of transcriptional variation has become an essential part of linking genotype to phenotype^42^. Despite the increasing scale of eQTL studies (e.g. 5,257 samples in Yao *et al*.^6^, and 2,116 in Zhernakova *et al*.^14^), fundamental questions about study design and analysis strategies have remained unanswered. Here, we have investigated the sensitivity and FDR of diverse multiple testing strategies, the factors contributing the identification of the causal eSNP, and the correction of eQTL effect size overestimation using a simple tool, BootstrapQTL. The insights from our simulation study are likely not limited to eQTL analysis, and may extend to other studies of genome-related quantitative traits, such as chromatin accessibility, methylation and other epigenetic traits.

## Methods

### Simulating genotypes and selecting eQTLs

Genotype data were simulated using HAPGEN2^31^ based on the FIN haplotypes of chromosome 22 from the 1000 Genomes Project data (phase3, GRCh37)^32^. The simulated genotypes had similar LD patterns with the reference data. Six sets of genotype data were generated at varying sample sizes: 100, 200, 500, 1000, 2000, and 5000 individuals. After filtering out SNPs with MAF less than 0.5% or Hardy-Weinberg Equilibrium (HWE) p-value less than 5×10^-6^, approximately 150 thousand SNPs remained in each data set.

We explored six different eSNP MAFs (0.5%, 1%, 5%, 10%, 25%, 50%) in each of the six genotype datasets, resulting in 36 scenarios in total. In each scenario, 200 SNPs at the scenario MAF were randomly chosen as true causal eSNPs, each regulating the expression of a randomly selected *cis* gene (within ±1 Mb from transcription start site of the gene). These 200 causal eSNPs were selected from an LD pruned subset where the pairwise r^2^ was ≤0.3.

### Simulating gene expression

To get a distribution of *cis*-eQTL effect sizes, we first performed eQTL mapping in DILGOM datase^33,34^ using additive linear model with covariates that accounted for gender, age, and population structure. Expression data were further scaled to make each gene’s expression across samples follow a standard normal distribution. To avoid an inflated number of associations due to LD structure among variants, we kept only the best association with the minimum nominal p-value for each gene. As shown in simulation results, only eQTLs with large effect sizes could be identified given a limited sample size. To reduce the bias caused by limited power, we included all genes to obtain the effect size distribution and fit it with a gamma distribution, from which we randomly selected true effect sizes.

In each scenario, 200 genes out of 618 genes on chromosome 22 were designated as “true eGenes” regulated by a causal eSNP each and the resting 418 as “null genes” with no truly associated eSNPs. The 200 true associations were modelled by a simple linear regression:

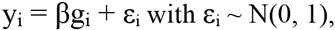

where y_i_ denoted the expression level of an eGene for individual i, β the genetic effect size of the corresponding eSNP, *g*_i_ the minor allele dosage of the eSNP coded as 0, 1 or 2, and ε_i_ the error variance for the i^th^ individual, which followed the standard normal distribution. For 418 null genes, no genetic effects were simulated (β = 0) and the simulated expression was normally distributed. True eGenes effect sizes were randomly drawn from a gamma distribution derived from a real dataset as described above.

### Mapping eQTLs and correcting for multiple testing

For *cis*-eQTL analysis, we used Matrix eQTL (version 2.1.1)^35^ to fit linear regression models between each gene and the minor allele dosage of all SNPs located within 1 Mb of their transcription start site. To adjust for multiple tests, we applied either (1) a correction method to all hypotheses (pooled method), or (2) a hierarchical correction procedure, where two methods were used in combination to correct for multiple SNPs tested for each gene and multiple genes separately.

Pooled multiple testing correction was performed using either Bonferroni correction or FDR-controlling procedures applied to all SNP–gene hypothesis tests. Bonferroni correction (pooled Bonferroni), Benjamini-Hochberg^17^ (pooled BH), and Benjamini-Yekutieli^18^ (pooled BY) FDR procedures were performed using “p.adjust” function in R (version 3.1.3)^43^, and Storey-Tibshirani^19^ (pooled ST) procedure was performed by the R package “qvalue” (version 1.43.0)^44^.

A three-step procedure was employed to perform hierarchical multiple testing correction. In Step 1, p-values of all *cis*-SNPs were adjusted for multiple testing for each gene separately (locally adjusted p-value). In Step 2, the minimum adjusted p-value from Step 1 was taken for each gene, then these adjusted p-values were further adjusted for multiple testing across all genes (globally adjusted p-value). Finally, in Step 3, significant eSNPs were identified for each significant eGene as SNPs with a locally adjusted p-value from Step 1 < the locally adjusted minimum p-value corresponding to the globally adjusted p-value threshold of 0.05.

Hierarchical multiple testing correction was performed using different combinations of multiple testing correction methods in Step 1 and Step 2 described above. In Step 1, we applied FDR procedures (ST, BH, or BY), Bonferroni, eigenMT^22^, or permutation approaches to correct for multiple local SNPs tested for each gene. In Step 2, we applied three FDR-controlling procedures, or Bonferroni correction to control the rate of false positive eGenes. Note that eigenMT and permutation approaches are used hierarchically by design.

When Bonferroni was used as a local correction method, the adjusted p-value was calculated by multiplying each linear model p-value by the number of SNPs in the corresponding 1Mb *cis* window for the tested gene. When using eigenMT, the linear model p-value was multiplied by the number of effective independent tests estimated from the genotype correlation matrix by eigenMT (in python 2.7.3)^22^. Permutations were performed by shuffling sample labels of expression data. For each gene, minimum nominal p-values from all permutation tests were kept to obtain the null distribution. Permutation p-values were calculated as the proportion of permutations showing more significant minimum p-value than the observed nominal p-value. The null distribution used to calculate permutation p-values was either (1) the exact distribution from permutations (exact permutation scheme), or (2) a beta distribution approximation of the null distribution tail, which is implemented in FastQTL (version 2.0)^20^. When using FastQTL, we performed either a fixed number of permutations (1,000), or under an adaptive scheme, a number ranging from 100 to 10,000 permutations determined via iterative estimates of gene significance throughout the permutation procedure.

### Correcting for Winner’s Curse

To evaluate and correct the effect of the Winner’s Curse, we considered the effect size estimates of the SNP with the minimum p-value (top eSNP) for each eGene. We use 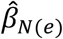 to denote the “naïve estimator”: the beta coefficient obtained from the linear regression of each eGene on its top eSNP.

We adjusted a bootstrap method^37^ to re-estimate eQTL effect sizes of significant eGenes determined by a hierarchical correction procedure (Bonferroni-BH by default; eigenMT-BH is also recommended). This approach consists of a repeated bootstrap analysis, in which random samples are drawn with replacement from the study dataset to partition the study samples into two groups: a bootstrap detection group of identical size to the original dataset comprising samples randomly selected with replacement, and a bootstrap estimation group comprising the remainder of the study samples. Due to the sampling with replacement, the bootstrap detection group typically comprised 63.2% of the study samples while the bootstrap estimation group comprised the other 36.8% of samples. The effect size is then estimated separately in the bootstrap detection and estimation groups for each eGenes and its top eSNP based on the original dataset.

After performing this bootstrap procedure with 200 bootstraps, three bootstrap estimators were calculated and compared for eGene effect size re-estimation: a shrinkage estimator:

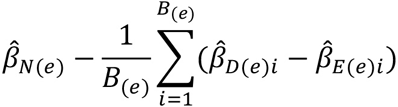

an out-of-sample estimator:

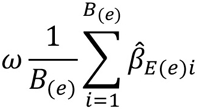

and a weighted estimator:

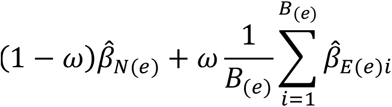

Where 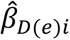 denotes the effect size of eGene *e* in each bootstrap detection group *i*, 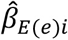 denotes the effect size of eGene *e* in each bootstrap estimation group *i*, and *B*_(*e*_) denotes the number of bootstraps in which the association between the eGene *e* and its top eSNP was significant in the bootstrap detection group (thus *B*_(*e*)_ ≤200). An association between an eGene and its top eSNP was considered significant in the bootstrap detection group if its locally adjusted p-value (corrected for multiple cis-SNPs within 1Mb of the respective eGene using e.g. eigenMT or Bonferroni) was smaller than the locally adjusted p-value corresponding to the 0.05 threshold after global adjustment (e.g. BH) in the eGene detection analysis prior to performing the bootstrap procedure. For the weighted estimator, the weight *w* was 0.632, *i.e.* the proportion of unique samples in the bootstrap detection group.

## Acknowledgements

This study was supported by the National Health and Medical Research Council (NHMRC) of Australia (grant no. 1062227) and the National Heart Foundation of Australia, and it was supported in part by the Victorian Government’s OIS Program. MI was supported by a Career Development Fellowship co-funded by the NHMRC and the National Heart Foundation of Australia (no. 1061435). We thank James E. Peters for helpful input and comments on the manuscript.

